# An array of *Zymoseptoria tritici* effectors suppress plant immune responses

**DOI:** 10.1101/2024.03.12.584321

**Authors:** E. Thynne, H. Ali, K. Seong, M. Abukhalaf, M. A. Guerreiro, V. M. Flores-Nunez, R. Hansen, A. Bergues, M. J. Salman, J. J. Rudd, K. Kanyuka, A. Tholey, K. V. Krasileva, G. J. Kettles, E. H. Stukenbrock

## Abstract

*Zymoseptoria tritici* is the most economically significant fungal pathogen of wheat in Europe. However, despite the importance of this pathogen, the molecular interactions between pathogen and host during infection are not well understood. Herein, we describe the use of two libraries of cloned *Z. tritici* effectors that were screened to identify effector candidates with putative pathogen associated molecular pattern (PAMP) triggered immunity (PTI)-suppressing activity. The effectors from each library were transiently expressed in *Nicotiana benthamiana*, and expressing leaves were treated with bacterial or fungal PAMPs to assess the effectors’ ability to suppress reactive oxygen species (ROS) production. From these screens, numerous effectors were identified with PTI-suppressing activity. In addition, some effectors were able to suppress cell death responses induced by other *Z. tritici* secreted proteins. We used structural prediction tools to predict the putative structures of all of the *Z. tritici* effectors, and used these predictions to examine whether there was enrichment of specific structural signatures among the PTI-suppressing effectors. From among the libraries, multiple members of the killer protein-like 4 (KP4) and killer protein-like 6 (KP6) effector families were identified as PTI-suppressors. This observation is intriguing, as these protein families were previously associated with antimicrobial activity rather than virulence or host manipulation. This data provides mechanistic insight into immune suppression by *Z. tritici* during infection, and suggests that similar to biotrophic pathogens, this fungus relies on a battery of secreted effectors to suppress host immunity during early phases of colonisation.

## Introduction

*Zymoseptoria tritici* is a major fungal pathogen of wheat, particularly in Europe, and is responsible for Septoria tritici blotch (STB) disease^1,2^. This fungus is unusual, in that it undergoes an extended latent asymptomatic growth phase which can last over two weeks under field conditions. During this phase, the fungus grows epiphytically on wheat leaf surfaces, before invading leaves through open stomata and growing through the apoplastic space of the mesophyll^3,4^. Throughout this phase, there is minimal activation of host defences. The fungus then transitions to necrotrophy, which is accompanied by the appearance of macroscopic disease symptoms and death of host cells. While the fungus is infecting the host, the host expresses membrane-associated receptors that monitor the apoplastic space for pathogen-associated molecular patterns (PAMPs), such as bacterial flg22 or fungal chitin, or specific effectors. Upon recognition of these foreign elements, the receptors signal for PAMP-triggered immunity (PTI) or effector-triggered immunity (ETI), respectively^5,6^. Accordingly, it is assumed that during the asymptomatic phase, *Z. tritici* secretes effectors into the apoplastic space to suppress PTI and ETI^7,8^.

Although hundreds of effector proteins have been predicted computationally from genome and transcriptome data^9–11^, only a few have been functionally characterised. The effectors *AvrStb6*, *AvrStb9* and *Avr3D1* have been shown to trigger ETI responses on the wheat with resistance genes *Stb6*, *Stb9* and *Stb7* respectively^12–14^. AvrStb9 contains a protease domain, and it is speculated that this domain contributes towards its virulence function. However, the functions of AvrStb6 and Avr3D1 have yet to be demonstrated. Another effector, ZtSSP2, has been demonstrated to interact with a wheat E3-ubiquitin ligase and this interaction is hypothesised to suppress PTI responses^15^, though this hypothesis remains to be conclusively proven. The most well-studied are the LysM domain-containing effectors, that sequester free chitin before it is recognized by the host, and offers a protective coat to hyphae from host secreted chitinases^7,16,17^. Via this mechanism, the pathogen can mask its own presence and evade host defences. However, *Z. tritici* mutants lacking LysM domain effectors remain partially virulent, suggesting the existence of other immune suppressing effectors produced by this fungus. No other *Z. tritici* effectors have been observed as active PTI suppressors.

High-throughput screening of fungal effectors in wheat still has technical difficulties, despite improvements in wheat protoplasts^18,19^ or via viral expression^20^. For ease of analysis, we chose to screen the effectors in the model organism *N. benthamiana.* Perception of PAMPs and apoplastic effectors often relies on activity of cell-surface receptor-like proteins (RLPs) or receptor-like kinases (RLKs). In many cases, receptors must partner with other cell-surface co-receptors such as BRASSINOSTEROID INSENSITIVE 1-associated receptor kinase 1 (BAK1) or Suppressor of BIR1-1/EVERSHED (SOBIR1/EVR) to initiate defence signalling^21^. We hypothesised that *Z. tritici* effectors that suppress conserved immune responses, such as BAK1-dependent responses, could be identified by screening their immune-suppressing activity in *Nicotiana benthamiana* (i.e., suppression of pathways conserved across monocots and dicots), and that these findings can be later translated into a wheat system. Herein we describe the independent screening of two different *Z. tritici* effector libraries, transiently expressed in *N. benthamiana,* to identify novel *Z. tritici* effectors with putative functions in suppression of PTI and ETI defence responses.

## Results

### Eleven candidate effectors selected as preliminary candidates for PTI suppression

In this study two libraries of effectors were examined, with different gene name identifiers^22,23^. The identifiers for each effector from both naming conventions (https://mycocosm.jgi.doe.gov^22^, <u>10.1534/g3.115.017731</u>^23^) are listed in *Supplementary File 1.* We first selected candidate effectors to establish our screen according to two main criteria: First, we considered that PTI-suppressing effectors show conservation among *Zymoseptoria spp.* as they are targeting core immune signalling processes. Second, we hypothesize that PTI-suppressing effectors will be specifically up-regulated during early plant colonization.

We first explored genomic data from five different *Zymoseptoria* species (*Z. tritici, Z. ardabiliae*^24^*, Z. brevis, Z. passerinii,* and *Z. pseudotritici*) to identify conserved orthologous effector candidates. Moreover, we included genome data from three *Z. tritici* isolates (Zt05, Zt09 (synonymous with IPO323), Zt10)^9,23^, considering that some effector genes can show presence-absence variation among individuals within the same species. We designed our analyses to identify orthologous genes present in all the analysed genomes. To this end, we performed an orthologue clustering analysis to identify shared effector orthogroups (1e-5 cut-off) resulting 56 orthogroups among the eight *Zymoseptoria* genomes (Supp. File 1).

Based on available RNA-seq data, we next selected *Z. tritici* orthologues from the 56 conserved orthogroups that were expressed during the symptomless growth phase^9^. Twenty-one effector candidates were highly expressed during the symptomless phase of infection (Table 1). Eleven candidates were most highly expressed during the necrotrophic phase, and thirty-four effectors displayed negligible expression during any phase of infection. We considered the 21 effectors as putative candidates that can suppress the PTI during the asymptomatic infection (Supp. File 1).

**Table 1:** Zt09 orthologues of effectors shared among all *Zymoseptoria spp.* that are highly expressed in either the necrotrophic or symptomless life-stages (FPKM values). Dark green = highest expression time-point, light green = second highest expression time-point. DPI: Days Post Infection). Gene models and accessions are from^23^ and FPKM values are from^9^.

| Effector library 1 |  |  |  |  |  |
| --- | --- | --- | --- | --- | --- |
| Effector ID | 3DPI | 7DPI | 13DPI | 20DPI | Predicted protein domain |
| Zt09_chr_5_00190 | 3 | 964 | 174 | 21 | No |
| Zt09_chr_1_01278 | 2.353 | 1,346.82 | 388.747 | 16.238 | No |
| Zt09_chr_1_02089 | 2,784.04 | 1,130.83 | 351.267 | 679.491 | No |
| Zt09_chr_1_01276 | 24.213 | 800.271 | 249.882 | 24.867 | No |
| Zt09_chr_4_00056 | 107.524 | 1,085.19 | 1,203.23 | 70.107 | No |
| Zt09_chr_4_00469 | 25.878 | 475.242 | 914.117 | 361.181 | No |
| Zt09_chr_8_00412 | 26.453 | 430.079 | 232.957 | 26.103 | LysM<sup>7,17</sup> |
| Zt09_chr_1_00132 | 0 | 330.243 | 115.889 | 6.076 | No |
| Zt09_chr_7_00276 | 425.071 | 695.194 | 602.686 | 596.913 | Cyclophilin-like |
| Zt09_chr_2_00242 | 43.248 | 219.234 | 116.208 | 39.932 | Hce2<sup>26</sup> |
| Zt09_chr_10_00356 | 123.602 | 124.803 | 86.445 | 109.563 | Duf2012 |
| Zt09_chr_3_00610 | 254.108 | 1,061.49 | 677.792 | 324.428 | Ribonuclease<sup>27</sup> |
| Zt09_chr_6_00044 | 390.542 | 215.367 | 101.428 | 65.288 | PR1-like |
| Zt09_chr_3_00904 | 7.177 | 3,079.04 | 1,092.28 | 77.793 | No |
| Zt09_chr_3_00971 | 8.88 | 1,138.54 | 623.621 | 18.977 | Arabfuran-catal |
| Zt09_chr_1_00805 | 0 | 43.218 | 3.891 | 0.784 | No |
| Zt09_chr_12_00080 | 192.981 | 223.174 | 193.916 | 175.779 | EMP24 |
| Zt09_chr_3_00667 | 13.644 | 96.777 | 343.599 | 23.132 | No |
| Zt09_chr_5_00497 | 192.017 | 175.528 | 80.582 | 120.353 | FAS1 |
| Zt09_chr_2_01151 | 35.845 | 19.719 | 984.238 | 328.263 | Cutinase |

We used InterproScan^25^ to add functional annotations to the 21 effector candidates. Ten effectors had predicted protein domains (not including HCE2 (an effector associated-domain, derived from *Cladosporium* Ecp2 effectors^26^); pfam: PF14856). Among these, we identified the previously characterized Zt6, a secreted ribonuclease with antimicrobial and cell-death inducing activity^27^, and also a LysM-domain containing effector underlining the suitability of our approach to identify functionally relevant genes (Table 1). Finally, we identified 11 symptomless phase-expressed effectors without known protein domains (with the exception of the HCE2 domain), and we focused our analyses on these unknown candidates.

### Five *Z. tritici* effector candidates suppress the flg22-induced PTI response

We then screened the 11 candidate effectors in *N. benthamiana,* to assess their ability to suppress a PAMP-triggered ROS burst, using the potent elicitor, flg22.

To establish an appropriate positive control for the assay, we surveyed orthologues of a known PTI-suppressing effector NIS1 identified in *Magnaporthe oryzae* (MoNIS1)^28^. Not only *Z. tritici* but also other *Zymoseptoria* sister species encoded orthologs of MoNIS1 (identified via BLASTp searches, Supp. File 1). In particular, *Z. tritici* IPO323 had two homologs, with only one expressed during the asymptomatic phase of infection (Supp. File. 1). We hypothesized that this protein (hereon described as ZtNIS1) would similarly inhibit PTI in *N. benthamiana*, akin to MoNIS1’s action.

In our transient gene expression assay, a control was expressed on the left half of the leaves, while comparison group was expressed on the right half to minimize biological variations that can arise from differences between and within leaves. The relative luminescence accumulation (RLU) for comparison groups was measured with respect to the control from the same leaf after flg22 treatments. We selected the hell-fire tag (HF tag), with an added fungal signal peptide, as our negative control. When the negative controls were expressed in both half of the leaves and PTI was induced with flg22, the RLUs was approximately one (Fig. 1), indicative of no PTI suppression. We then tested ZtNIS1 and MoNIS1 by transiently expressing each of these effectors on the right half of the leaves. The RLU for MoNIS1 and ZtNIS1 were significantly lower than the control experiment (Fig. 1), confirming that ZtNIS1 has similar PTI-suppressing activity as MoNIS1 and can serve as a positive control.

**Figure 1.**
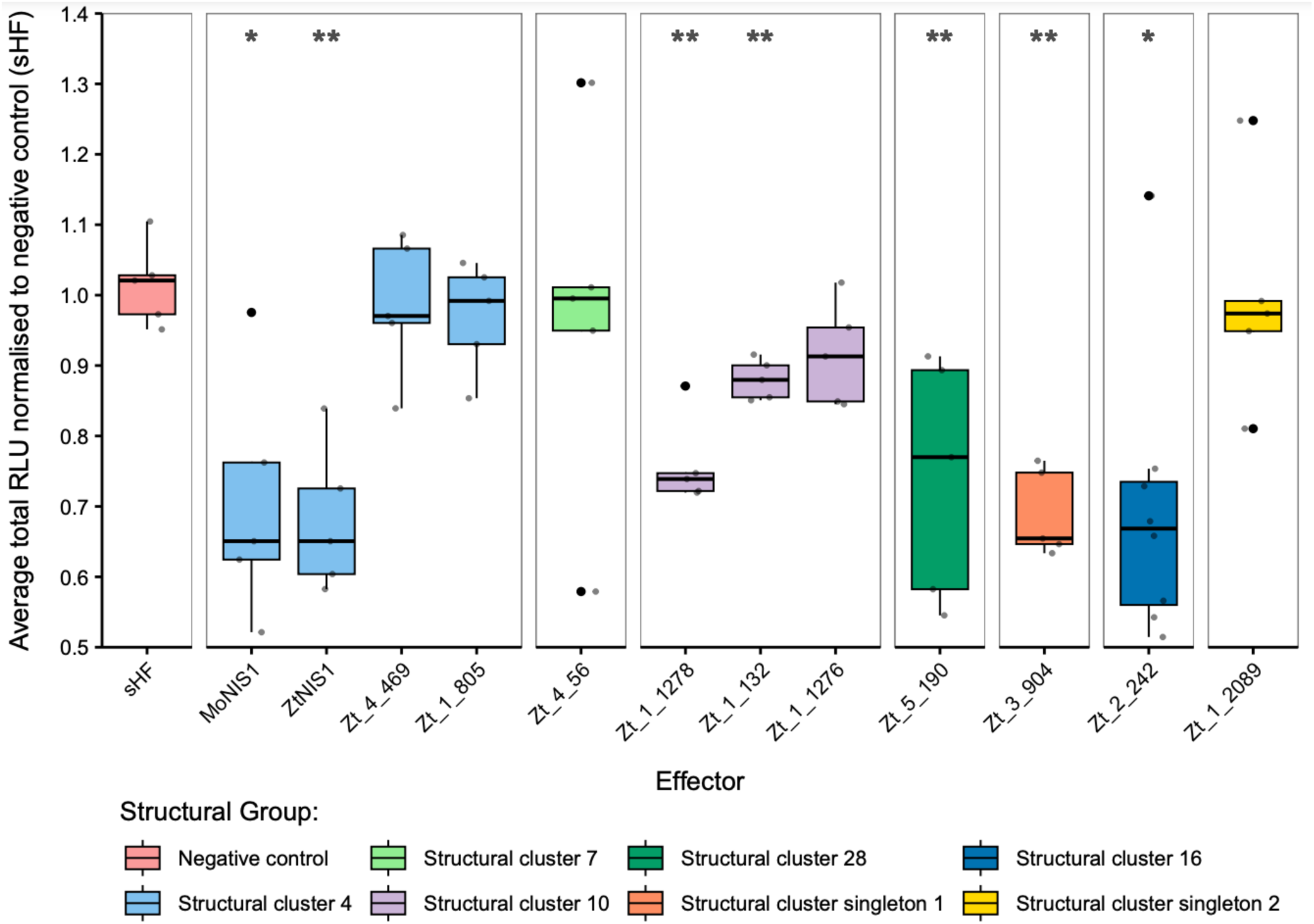
Various effectors from *Z. tritici* consistently suppress flg22-induced ROS burst. Candidate effectors were transiently expressed in *N. benthamiana*, with *Agrobacterium*. Each leaf had the negative control (sHF) expressed on one half, and an effector on the other. At 72 hours post infiltration (HPI), leaf discs from each side of a leaf were treated with flg22. The relative luminescence (RLU) from each ROS burst assay was measured by comparing the luminesnce of a comparison group to the negative control (sHF). Five effectors were identified as significant suppressors of flg22-induced ROS burst in comparison to the sHF controls (Wilcoxon test applied to assign significance).

Validating positive and negative controls in our assays, we assessed the suppressive ability of each of the 11 effectors on flg-22-induced ROS burst in *N. benthamiana* (Fig. 1). One of the effectors, Zt_3_00667, induced cell-death and was therefore excluded. Among the remaining ten effector candidates screened, five displayed significantly reduced RLU and were identified as putative suppressors of flg22-indcued ROS burst. These effectors were Zt_1_1278, Zt_1_132, Zt_5_190, Zt_3_904, and Zt_2_242. Among the observed immune-suppressors, Zt_1_132, displayed the weakest suppressive phenotype, with an average RLU of 0.88. The remaining suppressors have a greater magnitude of suppression, more similar to ZtNIS1.

### Additional *Z. tritici* candidate effectors suppress flg22-, β-glucan-, or chitin-triggered immunity when transiently expressed in *N. benthamiana*

Our initial screen indicated that five out of eleven tested candidate effectors suppressed the flg22-induced ROS burst. This relatively high incidence prompted us to question whether ROS burst suppression might be a common feature of *Z. tritici* effectors. To assess the prevalence of this phenomenon, we made use of an established library of cloned *Z. tritici* candidate effectors to uncover additional PTI-suppressing proteins. This second library contains 48 effectors that were identified as exhibiting elevated expression during the symptomless and transition phases of wheat leaf colonisation (Table 2)^10,27,29,30^. Each were previously cloned into *A. tumefaciens* expression vectors^29^. These effectors were not shown to induce cell-death in *N. benthamiana* and their virulence functions are currently unknown. These 48 candidate effectors were transiently expressed in *N. benthamiana* and tested for ability to suppress the ROS burst induced by either flg22 or the fungal PAMPs chitin and β-glucan (laminarin).

**Table 2:** Library of 48 *Z. tritici* effectors expressed in *N. benthamiana*^29^. Dark green = highest expression time-point, light green = second highest expression time-point. DPI: Days Post Infection). Gene models and accessions are from^22^ and FPKM values are from^10^.

| Effector Library 2 |  |  |  |  |  |  |
| --- | --- | --- | --- | --- | --- | --- |
| Effector ID | DPI1 | DPI4 | DPI9 | DPI14 | DPI21 | Predicted protein domain |
| 103555 | 71,8268 | 31,9067 | 1471,73 | 780,445 | 26,6313 | No |
| 106127 | 34,3027 | 38,5455 | 109,881 | 33,7765 | 21,0954 | No |
| 88698 | 252,613 | 270,297 | 741,601 | 84,5817 | 0,684321 | No |
| 67799 | 436,424 | 516,756 | 2230,69 | 281,55 | 4,51174 | No |
| 90776 | 44,4356 | 68,7214 | 1218,65 | 884,56 | 77,2719 | No |
| 103650 | 21,2892 | 38,4807 | 159,504 | 40,0327 | 12,5426 | FAS1 |
| 91702 | 4,10629 | 0 | 6,66709 | 91,1406 | 2,51597 | NTF2-like |
| 91885 | 6,55651 | 2,11865 | 31,1736 | 17,7543 | 3,20923 | No |
| 103900 | 133,184 | 394,016 | 4052,55 | 595,846 | 8,83538 | No |
| 92097 | 6,51416 | 16,9656 | 190,1 | 32,7757 | 1,60149 | Cellulase |
| 104000 | 439,149 | 534,497 | 871,931 | 493,021 | 24,8221 | No |
| 92792 | 5,27669 | 8,46092 | 13,2947 | 5,09436 | 1,90456 | No |
| 104404 | 498,735 | 496,239 | 3676,13 | 918,011 | 4,63377 | No |
| 93075 | 368,975 | 628,317 | 2148,16 | 116,477 | 4,20616 | No |
| 104794 | 1371,45 | 548,493 | 1995,37 | 2316,03 | 185,178 | No |
| 94107 | 36,7289 | 8,7981 | 75,3383 | 29,2692 | 3,53077 | No |
| 110052 | 10,2378 | 14,7326 | 75,8979 | 10,6483 | 1,14919 | No |
| 94290 | 0 | 18,6498 | 85,2821 | 7,00585 | 0,0334697 | No |
| 94526 | 110,308 | 218,86 | 1327,45 | 513,034 | 200,947 | No |
| 95478 | 11,674 | 11,1211 | 231,658 | 102,245 | 7,54892 | No |
| 105826 | 200,536 | 80,3845 | 178,797 | 66,0158 | 5,85966 | No |
| 96868 | 37,4388 | 31,5169 | 189,966 | 197,665 | 60,3986 | AltA1<sup>31</sup> |
| 106436 | 7,55897 | 238,087 | 478,146 | 17,871 | 0,663169 | No |
| 30802 | 68,6285 | 120,417 | 188,117 | 136,372 | 4,49828 | Metalloprotease |
| 34332 | 33,064 | 43,7393 | 42,5177 | 40,0953 | 49,4595 | Virginiamycin B lyase |
| 70022 | 8,06508 | 5,70359 | 5,81731 | 3,84823 | 2,4716 | No |
| 71681 | 36,6517 | 32,1224 | 93,2876 | 50,3864 | 10,7613 | Cellulase |
| 79286 | 24,3634 | 15,8572 | 33,5928 | 13,5946 | 13,91 | No |
| 82936 | 3,89403 | 4,66696 | 3,36315 | 5,41269 | 6,67506 | Cupredoxin |
| 89734 | 25,457 | 46,662 | 9,69304 | 7,20306 | 5,94365 | No |
| 91285 | 2,1966 | 0 | 0 | 3,9918 | 1,17446 | No |
| 91662 | 4,70605 | 315,802 | 1610,74 | 21,8738 | 0 | No |
| 94383 | 11,8254 | 52,3563 | 268,536 | 41,3623 | 15,9313 | No |
| 95416 | 282,183 | 135,761 | 513,047 | 98,1522 | 8,26741 | No |
| 95491 | 0 | 29,0063 | 7,67484 | 14,4753 | 197,868 | Hydrophobin |
| 95831 | 83,7147 | 22,9909 | 223,016 | 52,4644 | 9,66194 | No |
| 96389 | 57,4307 | 0 | 121,407 | 96,1556 | 8,39636 | No |
| 96543 | 147,111 | 722,166 | 79,0069 | 12,4891 | 6,65367 | Hydrophobin |
| 96865 | 2,1606 | 0 | 98,1958 | 42,4739 | 2,09637 | No |
| 97449 | 566,005 | 493,83 | 2595,6 | 300,374 | 1,07363 | No |
| 97526 | 0 | 60,2862 | 316,687 | 14,4184 | 1,21512 | No |
| 102996 | 75,3064 | 97,7416 | 16,2059 | 19,1364 | 104,235 | No |
| 104383 | 674,26 | 470,608 | 2172,05 | 287,264 | 28,7157 | No |
| 104444 | 146,918 | 1677,16 | 10414,2 | 1003,58 | 21,6612 | No |
| 105867 | 206,743 | 175,102 | 1068,21 | 244,413 | 28,646 | No |
| 105896 | 37,6148 | 25,0786 | 265,302 | 120,271 | 78,7094 | No |
| 107904 | 33,792 | 114,655 | 176,197 | 148,486 | 12,6837 | Hce2<sup>26</sup> |
| 111760 | 13,623 | 35,8904 | 189,232 | 29,0987 | 10,1225 | No |

In this screen, we used a secreted GFP (sGFP) as a negative control for ROS suppression and the *Pseudomonas syringae* effector, AvrPtoB (expressed intracellularly), was used as a positive control. In experiments with all three PAMPs, RLU values for AvrPtoB were consistently and significantly lower than the sGFP-expressing leaf discs indicating their suitability as controls. Suppression of the flg22-, laminarin, and chitin-induced ROS bursts were observed for nine, five and thirteen effectors respectively (Fig.2). In assays with flg22, the magnitude of ROS suppression by some *Z. tritici* effectors, whilst statistically significant, was weaker than that observed for AvrPtoB (Fig.2A). However, effectors 104404 and 104000 were notable as they suppressed ROS to a level similar to the AvrPtoB positive control. For laminarin-triggered ROS, we observed a suppressive phenotype for five effectors (Fig.2B). Similar to the flg22 assays, the suppressive effect caused by many effectors was less pronounced than by the AvrPtoB positive control, although still statistically significant. Only effector 104404 suppressed laminarin-induced ROS to a similar degree as AvrPtoB. For chitin-triggered ROS we found a suppressive phenotype for 13 effectors (Fig.2C). In contrast to the other PAMPs, the magnitude of ROS suppression following chitin treatment was often stronger, with several effectors exhibiting a potency similar to that of AvrPtoB. Across experiments, we found that twelve effectors suppressed the ROS burst for a single PAMP, three effectors suppressed ROS induced by two PAMPs, and three effectors suppressed the ROS induced by all three PAMPs tested. This data indicates that ROS suppression is a common feature shared by numerous *Z. tritici* candidate effector proteins.

**Figure 2.**
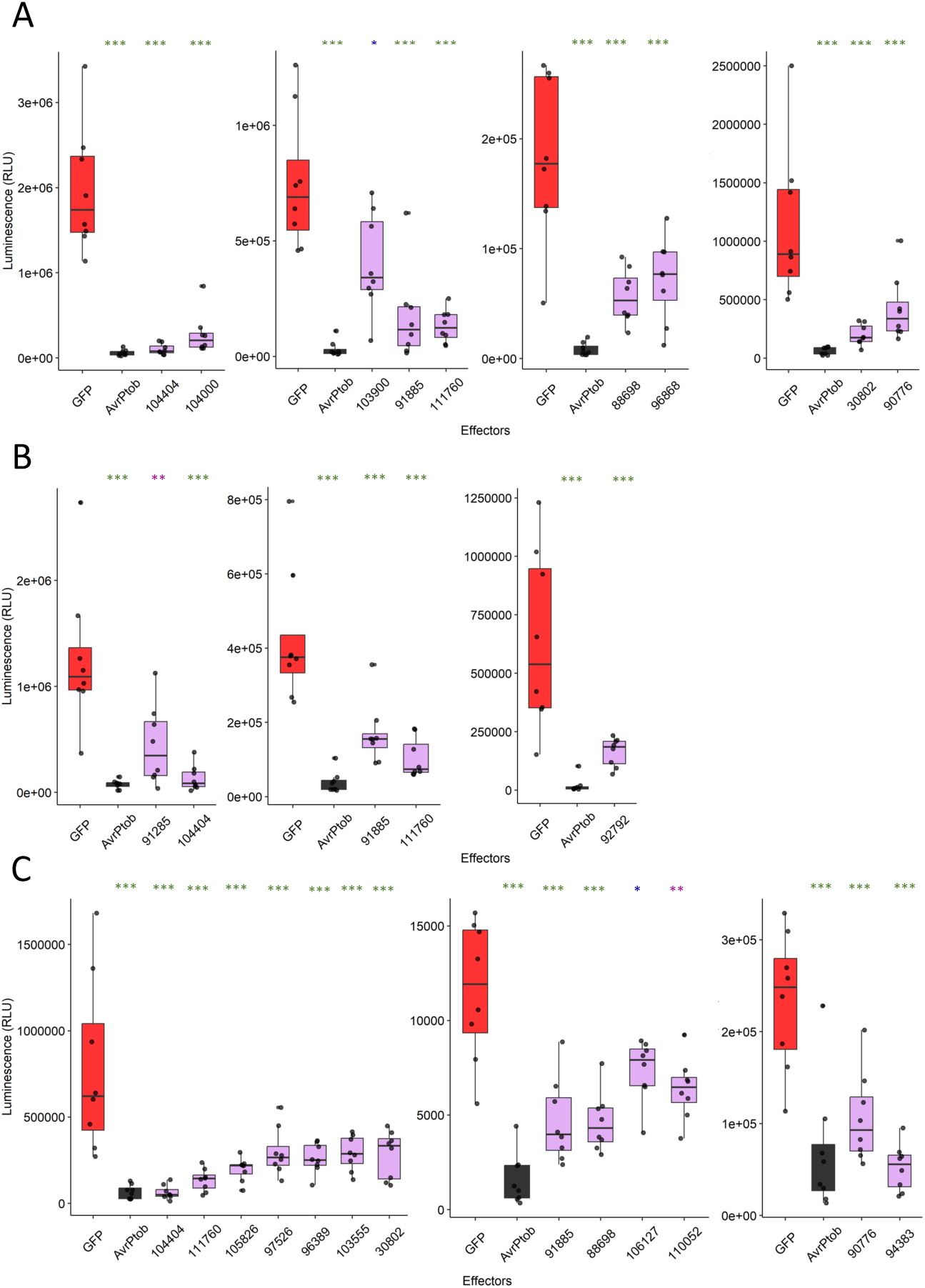
Suppression of the flg22-, laminarin- and chitin-induced ROS bursts. Candidate *Z. tritici* effectors were expressed in *N. benthamiana* and leaf squares used for ROS assay at 48 hpi. sGFP (shown in red) and AvrPtoB (shown in grey) were used as negative and positive controls for ROS burst suppression. A) flg22 treatment; B) Laminarin treatment; C) Chitin treatment. Asterisks indicate statistical significance at *p<0.05, **p<0.01, ***p<0.001 as performed by Tukey’s HSD test.

### Suppression of effector-induced cell death

We previously reported that several *Z. tritici* candidate effectors induce BAK1/SOBIR1-dependent cell death in *N. benthamiana*^29,30^. Given our recent observations of ROS burst suppression by a different subset of candidate effectors in the present study (Fig.2), we speculated that some of these proteins may have other immunosuppressive functions, including ability to suppress effector-triggered-immunity (ETI). To test this possibility, we co-expressed the cell death inducing effectors Zt6, Zt9, Zt11 and Zt12^27,29^ with the 48 candidate effectors described above (Fig.2). Co-infiltrations of cell death inducing proteins with sGFP were performed on the same leaves as controls. In these experiments, we observed repeated suppression of cell death by six candidate effectors (103900, 30802, 88698, 91885, 92097, 95478) (Fig.3). All six effectors were able to suppress Zt12-induced cell death, whilst three were also able to suppress Zt9-induced cell death. One effector, 92097, was able to suppress cell death induced by Zt9, Zt11 and Zt12. However, none of the effectors tested were able to suppress Zt6-induced cell death. This is consistent with Zt6 functioning as a ribonuclease toxin that initiates cell death independently of BAK1/SOBIR1^29^. Four of the six cell death-suppressing effectors were previously found to suppress ROS production induced by one or more PAMPs (Fig.2). This result indicates that *Z. tritici* candidate effectors are able to suppress multiple defence pathways thus contributing to evasion of immune surveillance.

**Figure 3.**
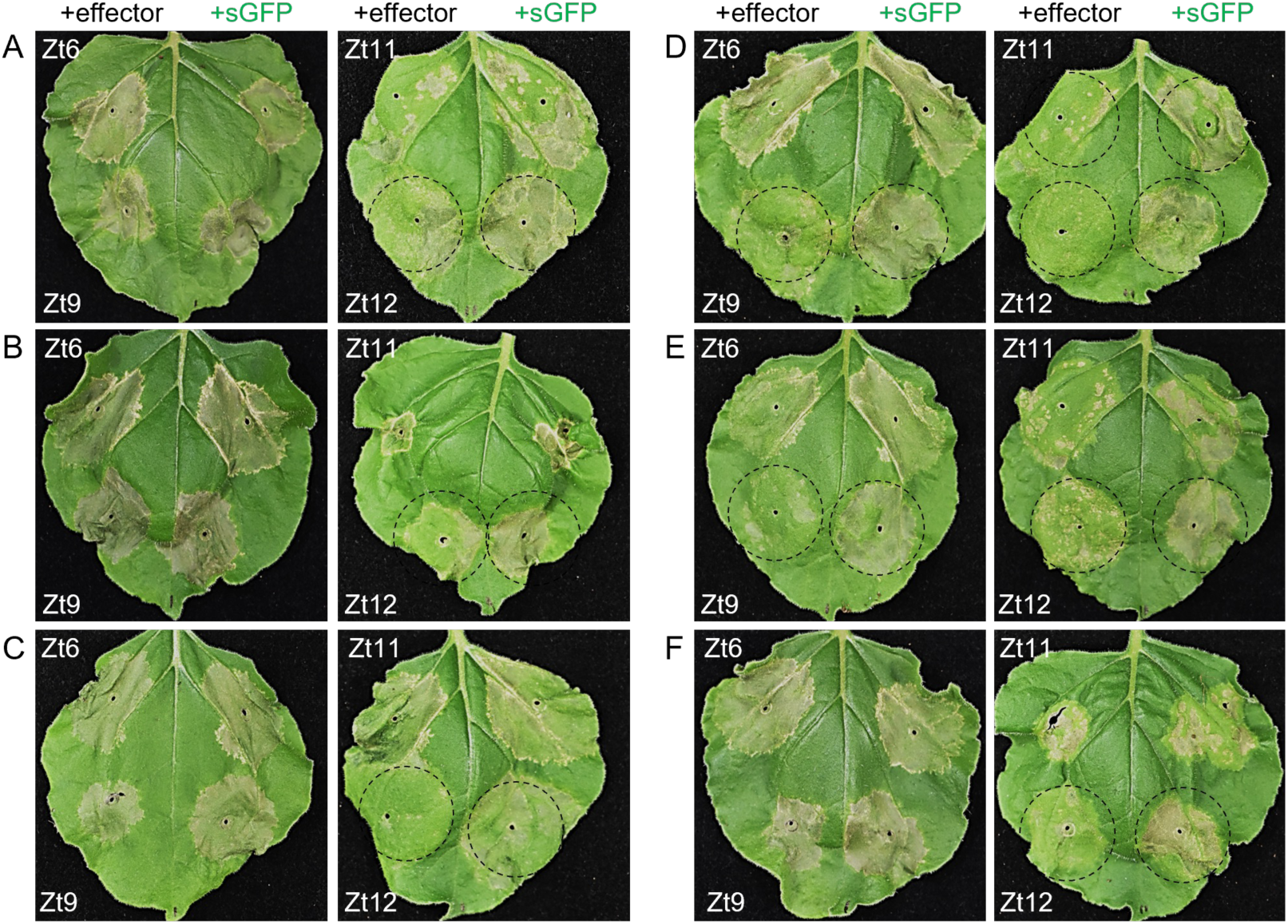
Suppression of effector-induced cell death in *N. benthamiana*. Leaves were co-infiltrated with *A. tumefaciens* strains delivering a cell death inducer (Zt6, Zt9, Zt11, Zt12) and either a negative control strain (+sGFP) or a candidate secreted effector (+effector). Effectors shown are (A) 103900, (B) 30802, (C) 88698, (D) 92097, (E) 95478, (F) 91885. Dashed circles indicate co-infiltration pairs where cell death suppression was observed in the effector treatment compared to the sGFP control. Leaves were photographed at 5 dpi.

### Structural predictions identified conserved folds among PTI-suppressing effectors

Structure prediction algorithms such as AlphaFold^32^ can offer novel insights into effectors that lack functional domains and sequence-related homologues. To identify possible commonalities among the PTI suppressors, we clustered whole proteome of *Z. tritici* IPO323 using structures predicted by AlphaFold^32^ (Supp. File 1). Where possible, we assigned the effectors of interest to specific structural families (Fig. 4; Supp. File 1). Among three effectors, 104404, 91885, and 111760 that suppressed the flg22-, laminarin- and chitin-induced ROS burst. 104404 was predicted to belong to a killer protein-like 4 (KP4-like fold) structural family. A reliable structure was predicted for 91885 with pTM score of 0.723, and it was clustered with two other effectors, 88619 and 106743, not tested in this study; however, no specific family was assigned to this cluster. In contrast, 111760 could not be accurately modelled. Three effectors, 88698, 30802, and 90776, suppressed both flg22- and chitin-induced ROS burst. The effector candidate 88698 belonged to the killer protein-like 6 (KP6-like fold) family, 30802 was predicted to be a metalloprotease based on structural similarity, and 90776 partially matched a pectate-lyase fold.

**Figure 4.**
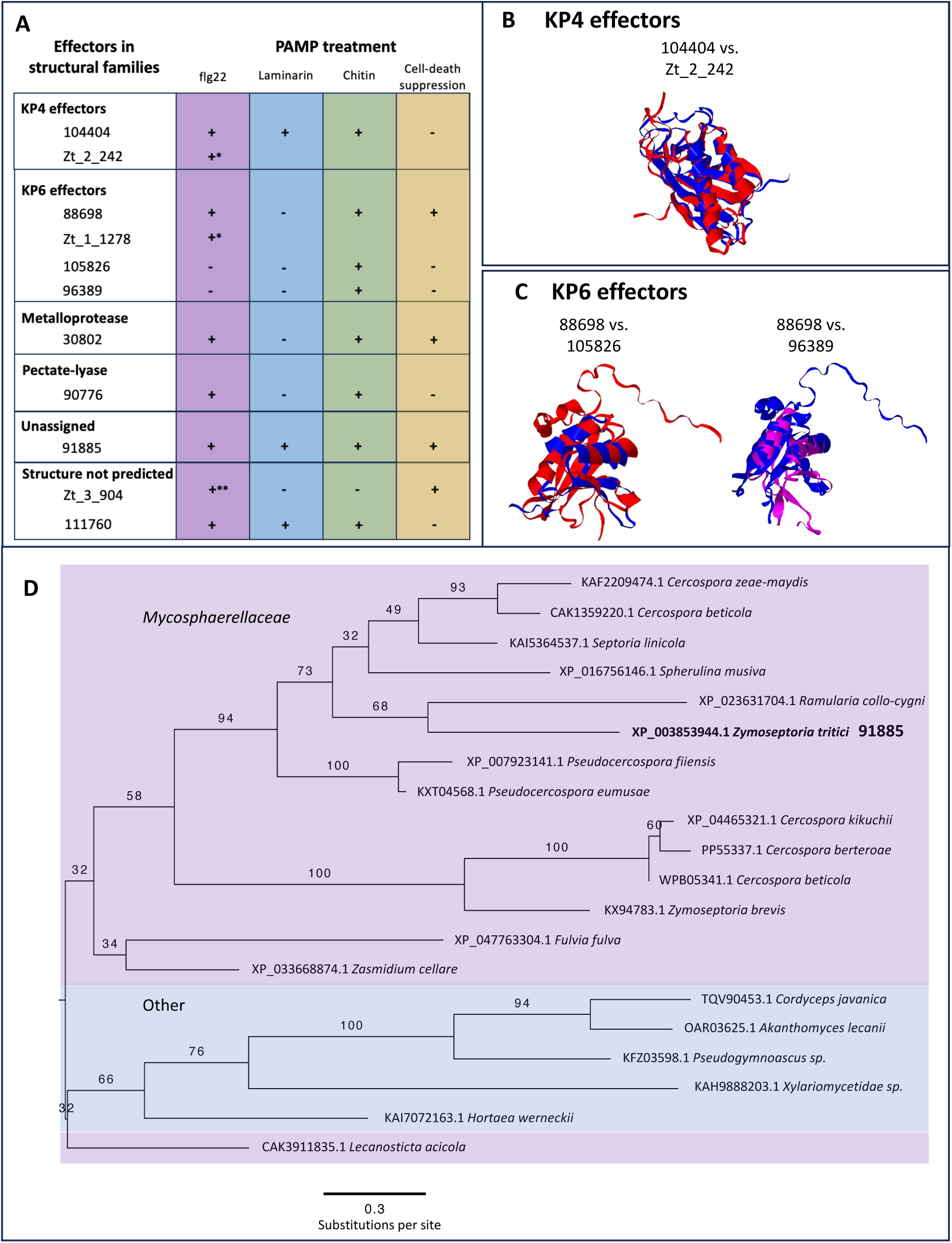
Multiple KP4-like fold and KP6-like fold effectors suppress PTI-responses A) Summary of selected effectors, their observed immune-suppressing activity, and predicted structural folds based on AlphaFold^32^. B) Structure alignment of the two immune-suppressing KP4-fold effectors (Red = 104404; Blue = Zt_2_242). C) Structural alignments of non-paralogous KP6-fold effectors. The structure of 88698 was used as the reference in both alignments (Red = 88698; Blue = 105826; Magenta = 96389. E) Phylogenetic tree of sequence homologues of the 91885 showing the occurrence of homologues across other fungal species.

In addition to 104404, another PTI-suppressing effector, Zt_2_242, was predicted to belong to the KP4 family. Despite sharing similar structures (Fig. 4B), these proteins do not share similarity at the sequence level. In total, seven effectors within the *Z. tritici* genome are predicted to belong to the KP4 family of effectors (Supp. File 1). Two more structural families were identified with multiple PTI-suppressing members. The first of these are the KP6-fold effectors, for which four were identified with varying PTI-suppressing activity. 88698 and Zt_1_1278 are paralogues, and both suppressed flg22-induced ROS burst. The other two KP6-fold effectors, 105826 and 96389, were not observed to suppress flg22-induced ROS burst. However, like 88698, they each suppressed chitin-induced ROS burst. 105826 and 96389 share no discernible sequence similarity with each other, or with either of 88698 or Zt_1_1278, but are similar in structure (Fig 4C). In total, *Z. tritici* is predicted to have nine KP6-fold effectors (Supp. File 1), which includes Zt9, previously found to trigger cell death in *N. benthamiana* and used as a treatment in the cell-death-suppression assay (Fig.3).

In addition to 111760, one other effector investigated here for which no specific 3D structure could be predicted, 103900, was identified as a PTI-suppressor. This effector is of interest as it was present in both libraries, and identified as a suppressor of flg22-induced ROS burst in both screens. The amino acid sequences of 111760 and 103900 were independently queried against the NCBI-NR database using BLASTp in order to identify sequence similar homologues. Homologues of 111760 were found among Mycosphaerellaceae species (Fig. 4D), whereas 103900 was limited to *Z. tritici* and some *Cercospora* species (Supp. File 1).

## Discussion

Despite the importance of *Z. tritici* as a major wheat pathogen, relatively little is known about the wheat-*Z. tritici* molecular interactions during the extended symptomless growth phase of infection. Our long-term goal is to identify and characterise effector proteins secreted during *Z. tritici* infection that target and suppress components of the wheat immune system, and in doing so, potentially identify host resistance or susceptibility factors. To support this goal, we established a high-throughput assay allowing us to screen multiple *Z. tritici* effector candidates with the overarching objective to identify immune-suppressing effectors. With our method based on heterologous expression, we were able to identify multiple *Z. tritici* effectors with PTI-suppressing activity. This greatly expands the number of effectors known to be functional during symptomless colonisation, beyond the previously described LysM-domain effector family^7,17^.

It is known that *Z. tritici* suppresses the wheat immune response during infection, and, further, *Z. tritici* infection can lead to systemic induced susceptibility (SIS), enabling non-adapted pathogens or avirulent isolates of *Z. tritici* to co-infect^33,34^. It is likely that SIS is induced as a result of effector manipulation of the host; for example, by altering long-range hormonal signalling. The receptors that monitor the apoplastic space, in which *Z. tritici* resides, can signal for changes in plant hormone and peptide signalling, altering the status of pathogen susceptibility^35–38^. Broadly, therefore, it is important that we study how pathogen effectors can be used to suppress or subvert receptor signalling. To this end, we first examined the function of ZtNIS1 to see if this *Z. tritici* effector displays similar BAK1-dependent immune-suppressing activity as described from orthologues in *Colletotrichum* and *Magnaporthe spp*^28^. Similar to the orthologues from these two species, the expressed *Z. tritici* homologue of NIS1 can suppress PTI responses. Our subsequent findings demonstrate that ZtNIS1 is not alone, and an array of *Z. tritici* effectors suppress plant immune responses.

Interestingly, we observed that some PTI-suppressing effector candidates share structural folds. The most represented was the KP6 fold, with four PTI-suppressing effectors. KP6-like effectors were first described from yeast, as virally encoded proteins with antimicrobial activity. They have subsequently been described from virus-associated maize fungal pathogen, *Ustilago maydis,* with antifungal activity^39^. A variety of structural prediction screens of plant pathogens found this fold to be well-represented^28,40–42^, and so, combined with our new data, there is evidence of this fold playing a role in plant-pathogen interactions. Despite our observations of these four *Z. tritici* KP6-fold effectors suppressing PTI, not all members in this structural family do. For example, one of the KP6-fold effectors, Zt9, is known to induce cell-death in *N. benthamiana* rather than to suppress immunity; however, this phenotype in *N. benthamiana* does not mean it is not a suppressor in wheat. This effector is one of the nine *Z. tritici* KP6-fold effectors, demonstrating potential variation in activity. KP6-fold effectors from *Cladosporium fulvum* have been screened in wild tomatoes and cell-death was observed. It is, therefore, possible that there are solanaceous receptors that recognise members of this structural family^43^. Follow-up analyses should investigate each of these nine *Z. tritici* homologues and determine which are PTI-suppressors, which induce cell-death, and what is the difference between each that results in these polarized phenotypes.

Surprisingly, two effectors from killer protein-like 4 (KP4) family were also identified with PTI-suppressing activity. KP4-fold effectors have also been described as antimicrobial having a calcium channel-inhibiting activity, when screened against mammalian, fungal, and plant cells^39^. During PTI, apoplastic calcium is an important signalling molecule and transported into the cell^44,45^. There is a close association between this PTI calcium signalling and other signalling responses, such as ROS burst^45^. Therefore, in the cases of *the Z. tritici* KP4-fold effectors, it is quite possible they are attenuating calcium signalling which in turn results in the observed ROS burst-suppression activity. Previously, a *Z. tritici* KP4-fold effector was identified after as a candidate necrosis-inducing effector (necrosis-inducing protein 2 (ZtNIP2; not screened in this study) from culture filtrate of the fungus^46^. Four *Fusarium graminearum* KP4-fold effectors have been described with putative roles in virulence in wheat^47^. Three of these *F. graminearum* effectors were identified in a cluster, and when the entire cluster was knocked-out, virulence in wheat seedlings was reduced and root development inhibited^47^. These previous findings, combined with our own, indicate a potentially important role for KP4-fold effectors in plant infection (aside from niche competition between fungi and other microbes).

Although we have chosen to highlight effectors belonging to specific and enriched effector fold families, multiple effectors were identified with putative immune suppressing activity from among our two libraries. For example, 91885, displayed PTI-suppressing activity for all treated PAMPs in our assays, and appears to be a conserved effector among the Mycosphaerellaceae (and clustered with two other *Z. tritici* effectors, 88619 and 106743). These effectors identified are all interesting candidates for downstream functional analyses, and their unscreened structural homologues should be examined for whether they possess similar immune-suppressing activity. However, we should emphasise that effector structural predictions are very useful for hypothesis generation, but should not be used to conclude specific function without validation. It should also be noted that our findings were obtained via screeing in *N. benthamiana. N. benthamiana* is a useful model for studying effector function due to ease of use for both *Agrobacterium* infiltration and testing immune responses. Hereby, several new studies have demonstrated the use of heterologous expression to characterize the role of plant pathogen effectors from *Z. tritici* and related species^48, 49^. However, this is still a non-host system, and the activity of the *Z. tritici* effectors identified here, should ideally be corroborated in wheat protoplasts or with viral expression in whole wheat plants.

Our findings suggest that immune suppression during the symptomless infection stage is an important part of colonisation. This is a relatively cryptic stage of growth and there is no evidence of *Z. tritici* feeding^4,20^. This emphasises the importance of the symptomless phase, developmentally, for the fungus and, accordingly, the importance of evading the host immune system. Most of the effectors examined in this study are primarily expressed during the symptomless phase; however, host recognition can occur earlier, during initial stomatal penetration. The avirulence effector, *AvrStb6*, is expressed during stomatal penetration. In wheat cultivars with AvrStb6’s corresponding resistance receptor, Stb6, infection is hindered at this early stage when the fungus grows through the stomatal opening^50,51^. A similarly timed phenotype is observed for another resistance to *Z. tritici* receptor, Stb16q^52^. This all occurs before expression of the immune suppressing effectors identified in this study, are at their peak. It is relevant to note that infection of a virulent strain of *Z. tritici* can enable subsequent infection of an independently avirulent strain, by inducing SIS^33,34^. Therefore, it is quite possible that timing of immune suppressing effectors plays an important role in SIS development, and inhibition of resistance gene function.

## Materials and methods

### Selection of candidate effectors

Candidate gene sets were selected and defined in two independent ways. Firstly, to conduct an initial screen we selected candidate genes according to expression pattern and sequence conservation across different *Zymoseptoria* species. Total protein sets were obtained for the three *Z. tritici* isolates (Zt05, Zt09, Zt10)^9^. Predicted proteins of *Z. ardabiliae* (Za17)^24^*, Z. pseudotritici* (Zp13)^24^, and *Z. brevis* (Zb18110)^23^ were obtained from the JGI Mycosm portal (Za17: https://mycocosm.jgi.doe.gov/Zymar1/Zymar1.home.html; Zp13: https://mycocosm.jgi.doe.gov/Zymps1/Zymps1.home.html; Zb18110: https://mycocosm.jgi.doe.gov/Zymbr1/Zymbr1.home.html). The protein set for *Z. passerini* was derived from the annotation presented in^53^.

Effectors from each protein set were predicted with the use of SignalP (v5.0b)^54^ and EffectorP (v2.0)^55^. Fasta files for predicted effectors are stored in the Zenodo page associated with this project (DOI: **10.5281/zenodo.10037259**). OrthoMCL predictions were performed with default settings (e-value -0.5). Input effector fasta files with edited names compatible with OrthoMCL and the OrthoMCL output files are deposited in the same Zenodo page (DOI: **10.5281/zenodo.10037259**). Effector gene expression for early colonization (three days post infection (3DPI)), asymptomatic growth (7DPI and 13DPI), and necrotrophic phase (20DPI), was obtained from the data set generated in^9^. Candidate effector expression levels were examined for the reference strain, Zt09 (synonymous with IPO323). All of the effector candidates and corresponding annotations are listed in *Supp. File 1* (including amino acid sequences)

Effector protein sequences were analysed with the InteproScan Geneious plugin (v.2)^25^ to predict protein domains. Similarly, phylogentic analyses were performed using the RAxML Geneious plugin (v.4)^56^, with a parsimony random seed value of 1,234, and 100 bootstrap replicates.

#### Clustering predicted structures

We aim to cluster the predicted structures of the whole proteome of *Z. tritici* IOP323. 10,689 predicted structure of *Z. tritici* IOP323 (taxonomy ID: 336722) were downloaded from the AlphaFold Database (Varadi et al., 2022). The structures of 992 secreted proteins were obtained from the previous study and replaced the models from the AlphaFold database if their averaged pLDDT scores were higher than the database structures^57^. This corresponded to 849 structures. The structures of three proteins (Zt_1_805, Zt_1_1278, and Zt_9_367), missing in *Z. tritici* IOP323, were predicted with AlphaFold v2.3.2 and included^32^.

Signal peptides predicted from SignalP v5.0^54^ were removed from the database structures. Low-confidence N- and C-terminal flexible stretches were trimmed off by examining the average pLDDT with a sliding window of four and a cutoff of 40. If the length and the average pLDDT scores of the remaining protein sequences were smaller than 50 amino acids or less than 60, respectively, the structures were discarded. The remaining 8,335 structures were clustered with FoldSeek (easy-cluster -s 7.5 -c 0.4 –alignment-type 1 –tmscore-threshold 0.5)^58^. This clustering output was compared to the one from the previous study^57^.

### Candidate effector synthesis and cloning

Full-length effector DNA sequences (intronless) and Zt_13_171 signal peptide for entry into destination vector to create the secreted tag)) were synthesized as gene fragments by TWIST Biosciences. Sequences were codon optimized for *N. benthamiana* expression, and synthesized with sequence overhangs compatible with BsaI cloning into the final vector plasmids (Effector sequences, with BsaI compatible overhangs for entry into the vector plasmid via GoldenGate cloning listed in (Supp. Table 2). The vector plasmid, pICSL22011 (with his/flag “hellfire” tag (HF tag) was kindly provided by Mark Youles (Synbio, TSL, Norwich, UK). Sequences were cloned into the vectors using the one-pot GoldenGate cloning method, using BsaI. Cloning product was transformed via heat-shock into chemically-competent Top10 *E. coli* cells for plasmid propagation. Plasmid inserts were Sanger sequenced by Eurofins Genomics (Ebersgerg, Germany), using primers from outside the insert site (Supp. Table 2).

### Transient expression assays in *Nicotiana benthamiana*

Plasmids generated for the construction of either effectors or control sequence (secreted hell-fire tag (sHF)), were transformed into *Agrobacterium tumefaciens* strain GV3101 and grown on solid DYT medium (Kanamycin (K), Gentamicin (G), and rifampicin (R) selection) at 28°C for two days. Single colonies were selected, and grown in liquid DYT (K+G+R selection) overnight, at 200RMP, at 28°C. Glycerol stocks were made from these cultures and stored at -80°C. Before *N. benthamiana* transformation, bacterial glycerol stocks were plated onto DYT (K+G+R selection) overnight at 28°C. *Agrobacterium* was scraped from plate into infiltration buffer (IB: MiliQ water, 10mM MgCl_2_-MES, acetosyringone), and incubated at room temperature for one hour. The OD600 was measured after one hour, and diluted in IB to a final OD600 of 0.5 (except of p19 silencing suppressor (kindly provided by M. Sauter, CAU, Kiel) which was included in every assay sample, at an OD600 of 0.1). *Agrobacterium* was infiltrated into four-to-five-week-old *N. benthamiana* leaves using a needleless 1ml syringe. For experiments performed at the University of Birmingham (Figs.2-3), *A. tumefaciens* GV3101 strains harbouring pEAQ-HT-DEST3 (effector) have been described previously. The pEAQ-HT-DEST3 (sGFP) strain was generated in this study by generating a pEAQ-HT-DEST3 construct harbouring the *N. tabacum* PR1a signal peptide (SP) fused to GFP. For ROS burst assays, all Agrobacterium strains were syringe infiltrated into leaves of 4-5 week-old plants at an OD_600_=1.2. For cell death suppression assays, all strains were prepared to an OD_600_=1.8 and mixed in a 1:1 ratio such that the final concentration of elicitor and sGFP/effector was OD_600_=0.9. Each experiment was performed thrice. (sGFP) or pEAQ-HT-DEST3 (AvrPtoB) were infiltrated into leaves at a final OD_600_=1.2.

### Elicitor-induced ROS burst suppression assays

For the initial method development, we used four-to-five-week-old *N. benthamiana* leaves. These were infiltrated with *Agrobacterium tumefaciens* (one half of leaf expressing sHF and the other half an effector candidate). Three days post infiltration, 36 leaf discs were harvested from each side of the leaf and placed in a white-bottomed 96-well plate (sHF leaf discs were placed in wells in rows A, C, and E, and effector leaf discs were placed in rows B, D, and F), in 200ul of MiliQ water. The plates were placed in the dark until use (six-to-nine hours). 20-40 mins before measurements, the 200ul of MiliQ water was replaced with 100ul of MiliQ water. Just prior to reading, leaf discs in rows 11 and 12 were treated with mock (20μM luminol and 1μg horse-radish peroxidase (HRP)) and leaf discs in rows 1-to-10 were treated with flg22 (12.5nM flg22, 20μM luminol and 1μg HRP, final concetration). Resulting RLU was measured over 30 minutes in a Tecan 200 Pro plate reader (Tecan, Männedorf, Switzerland) Temperature ranges of the plate reader used in these assays were from 20-to-26°C (below 20°C the ROS burst was reduced and above 26°C the ROS burst values were inconsistent, with leaf discs ranging from highly active to non-responsive).

For the screening of 48 additional effector candidates at the University of Birmingham, the following methods were used for elicitor treatments. Leaf squares approximately 3mm x 3mm were harvested from 5-week-old *N. benthamiana* plants with a scalpel and added to wells of 96-well plates containing 200μl dH_2_O. Plates were incubated in the dark overnight prior to performing the ROS burst assay. The following day, the dH_2_O in each well was removed immediately prior to the assay, and replaced with 150 µl of assay solution containing HRP (20ng/ml), luminol L-012 (20μM) and either flg22 (100nM), chitin (100μg/ml) or laminarin (100μg/ml). Luminescence was captured over 2 hours (90 cycles) using a PHERAstar FS plate reader (BMG Labtech) controlled through the PHERAstar control software. Each plate contained eight replicates of each effector or control treatment. These experiments were repeated thrice.

### Data availability

Datasets (predicted effector sets, OrthoMCL output data, and raw and curated ROS burst data sets, structural prediction data) have been uploaded to the project’s Zenodo page (DOI: **10.5281/zenodo.10037259**) and/or in Supp. File 1. Within this Zenodo page we have also included IP/MS data for ZtNIS1, performed in *N. benthamiana*, identifying putative interaction partners. All plasmids (effector *N. benthamiana* expression plasmids and Y2H plasmids) are available upon request (for material transfer agreements relating to use of pICSL22011 plasmids, please contact Mark Youles, SynBio, The Sainsbury Laboratory, Norwich, U.K.). Use of the pEAQ-HT-DEST vector system is done so under license from Plant Bioscience Ltd/Leaf Systems International Ltd (Norwich, UK) to Graeme Kettles (University of Birmingham).

## Acknowledgements

The authors thank Mark Youles for providing the plasmids pICSL22011 (HF-tag) and pICSL22012 (GFP-tag). GK and HA are grateful to Luke Alderwick for guidance on use of the PHERAstar plate reader.

## Funding

E. Thynne was funded by a Marie Skłodowska-Curie Early-Stage grant from the European Research Commission. H. Ali was supported by a PhD scholarship awarded by the Darwin Trust of Edinburgh. Rothamsted Research receives strategic funding from the Biotechnology and Biological Sciences Research Council of the United Kingdom (BBSRC). This work was further supported by the European Research Council under the European Union’s Horizon 2020 research and innovation program (consolidator grant FungalSecrets, ID 101087809 to E. Stukenbrock). We acknowledge support from the Delivering Sustainable Wheat [BB/X011003/1] and Growing Health Institute Strategic Programmes [BB/X010953/1; BBS/E/RH/230003A].

## Supplementary Material

Supplementary_File_1

## Notes

### Competing Interest Statement

The authors have declared no competing interest.

